# Anthropogenic factors influence the occupancy of an invasive carnivore in a suburban preserve system

**DOI:** 10.1101/2020.02.10.939959

**Authors:** John P. Vanek, Andrew U. Rutter, Timothy S. Preuss, Holly P. Jones, Gary A. Glowacki

## Abstract

Domestic cats (*Felis catus*) are one of the world’s most damaging invasive species. Free-ranging cats kill billions of wild animals every year, spread parasites and diseases to both wildlife and humans, and are responsible for the extinction or extirpation of at least 63 species. While the ecology and conservation implications of free-ranging cats have well studied in some locations, relatively little is known about cats inhabiting urban nature preserves in the United States. To address this knowledge gap, we used camera traps to study the occupancy and activity patterns of free-ranging cats in 55 suburban nature preserves in the Chicago, IL metropolitan area. From 2010–2018 (4,440 trap days), we recorded 355 photos of free-ranging cats across 26 preserves (*ψ*_naïve_ = 0.45) and 41 randomly distributed monitoring points (*ψ*_naïve_ = 0.18). Cats were detected every year, but rarely at the same point or preserve, and cats were largely crepuscular/diurnal. Using single-season occupancy models and a “stacked” design, we found that cat occupancy increased with building density and detectability was highest near the urban/preserve boundary. Based on our top-ranked model, predicted occupancy within individual preserves ranged from 0.09 to 0.28 (*ψ*_mean_ = 0.11) and was poorly correlated with preserve size or shape. Overall, our results suggest that free-ranging cats are rare within suburban preserves in our study area, and that these cats are most likely owned or heavily subsidized by people (which pose different risks and management challenges than truly feral cats). We discuss the conservation and management implications for urban natural areas.

**Highlights:**

- We surveyed for domestic cats across 55 suburban preserves from 2010-2018.
- We modeled occupancy and detectability as a function of urban covariates.
- Cat occupancy was low overall and best predicted by building density.
- The risk to native species is highest near preserve boundaries bordered by built environments.

## 1. Introduction

Following a global trend of urbanization (Vitousek et al. 1997), most people in the United States now live in urban areas (U.S. Department of Housing and Urban Development and U.S. Census Department 2017). However, the area occupied by urban landcover (indicative of suburban development), has greatly outpaced the rate of urban population growth itself (Heimlich and Anderson 2001, Destefano et al. 2005). Suburban areas differ from that of intensely urban city centers (e.g. Manhattan or downtown Chicago) and typically consist of single-family homes (single or double-storied) with lawns and backyards spaced at a moderate to high density, interspersed with light industry, basic services, and multi-family homes (Marzluff et al. 2001). This suburban expansion or “urban sprawl” is a major threat to biodiversity, as it spreads the most pernicious threats of urbanization (i.e. habitat loss, habitat fragmentation, and invasive species) outward from city centers into the surrounding landscape (Czech et al. 2000, Marzluff et al. 2001, Marzluff 2002, McKinney 2002, 2008).

Domestic cats (*Felis catus*) are an invasive species common in urban and suburban areas. Free-ranging domestic cats (i.e. cats that are outside direct supervision of a human including pet cats that are allowed outside, barn cats, “community” cats, and feral or wild cats) are one the world’s most damaging invasive species (Lowe et al. 2000, Doherty et al. 2016). Free-ranging cats kill billions of birds, mammals, and reptiles every year (Blancher 2013, Loss et al. 2013, Woinarski et al. 2017, 2018). Cats also spread parasites and diseases to both humans and wildlife (Dubey and Jones 2008, Gerhold and Jessup 2013, Ma et al. 2018), and are responsible for the extinction or extirpation of at least 63 species (reviewed in Doherty et al. 2016). For example, cat depredation led to the extinction of the Stephens Island Wren (*Traversia lyalli*) (Galbreath and Brown 2004) and the extirpation of the Estanque Island population of the Ángel de la Guarda Deer Mouse (*Peromyscus guardia*) (Vázquez-Domínguez et al. 2004).

The ecological impacts of free-ranging cats are not limited to islands or wilderness areas. Cats in urban areas prey on native species and can occur at densities much greater than that of native carnivores (Churcher and Lawton 1987, Coleman and Temple 1993, Burton and Doblar 2004, Lepczyk et al. 2004). For example, Balogh et al. (2011) reported that domestic cats were responsible for 47% of known predation events on radio-tracked fledglings, and Flockhart et al. (2016) estimated a free-ranging cat density of up to 49 cats/ha in Guelph, Canada. Cats in urban areas tend to have smaller home ranges than those in rural or wild landscapes (Horn et al. 2011, Hall et al. 2016, Hanmer et al. 2017), but they still venture into urban greenspaces, such as parks and nature preserves (VanDruff and Rowse 1986, Kays and DeWan 2004, Morgan et al. 2009, Wierzbowska et al. 2012, Gehrt et al. 2013), where potential impacts on biodiversity are likely larger. Habitat selection of free-ranging cats, particularly in urban areas, is highly variable and likely location specific. Studies from New Zealand to Illinois, USA have reported selection for both urban and natural areas, avoidance of natural areas, and no habitat selection (Metsers et al. 2010, Horn et al. 2011, Gehrt et al. 2013, Kays et al. 2015, Elizondo and Loss 2016). Clearly, more research is necessary to better explain the habitat selection and space use of free-ranging cats in urban areas.

In this paper, we describe the spatial ecology of free-ranging cats inhabiting 55 nature preserves in the suburbs of the third largest metropolitan area in the United States: Chicago. While the demography and movements of free-ranging cats have been studied extensively in dense urban areas, relatively few studies explore the ecology of cats inhabiting suburban nature preserves, and fewer still take a landscape-scale approach (Kays and DeWan 2004, Morgan et al. 2009, Kays et al. 2015). Further, while anthropogenic factors have been linked to cat population parameters (e.g. Flockhart et al. 2016), few studies have compared the relative utility of different urban metrics (e.g. building density versus percent impervious surfaces) as predictors of population parameters. To address this deficit, we analyze nine years of systematic camera trapping data from a large-scale and on-going suburban wildlife monitoring program. Specifically, we (1) use occupancy modeling to compare the relative importance of different urban covariates, (2) explore temporal patterns of cat activity, and (3) examine the relationship between preserve design (e.g. size, shape, degree of urbanization) and cat occupancy.

To the best of our knowledge, this is the first study to generate detection-corrected, spatially explicit estimates of free-ranging cat occupancy across a landscape of suburban preserves. The results from this study can be used by land managers, conservation biologists, and urban planners to aid in the management of free-ranging cats, to help develop conservation plans for cat-sensitive species, and to guide the design of suburban nature preserves.

## 2. Methods

### 2.1. Study Location

Our study took place in Lake County, IL (land area = ∼1150 km^2^). Lake County is a highly urbanized suburb in the Chicago Metropolitan Area (Figure 1). Lake County is one of the most densely populated counties in the United States with ∼700,000 people and a population density of 607 persons/km^2^ (United States Census Bureau 2018), and the greater metro area has >10,000,000 inhabitants. Prior to European settlement (pre 1830), Lake County was a mosaic of savanna (45%), prairie (30%), and woodland/forest (15%) (Bowles and McBride 2005), but today is dominated by anthropogenic features (Figure 1). Within this urban landscape, the Lake County Forest Preserve District (LCFPD) manages 55 “forest preserves”, totaling 120 km^2^ for multiple uses, including biodiversity conservation and outdoor recreation. During our study, dominant plant communities within LCFPD preserves included forest (28%), wetlands (17%), and old fields (15%). Historically dominant communities such as prairie and savanna were uncommon (8% and 5%, respectively). Developed land (including turf grasses) was rare (3%) within the preserve boundaries themselves, although this excludes public roads and private inholdings. Other community types (e.g. crops, woody shrubs) make up the remaining 24% (XX, unpublished data). The climate in Lake County is temperate with precipitation averaging 93 cm/year (Illinois State Climatologist 2019). In this paper we refer to these “forest” preserves as “preserves” or “nature preserves,” but point out we are not referring to Illinois Nature Preserves as designated by the Illinois Nature Preserve Commission.

**Figure 1.**
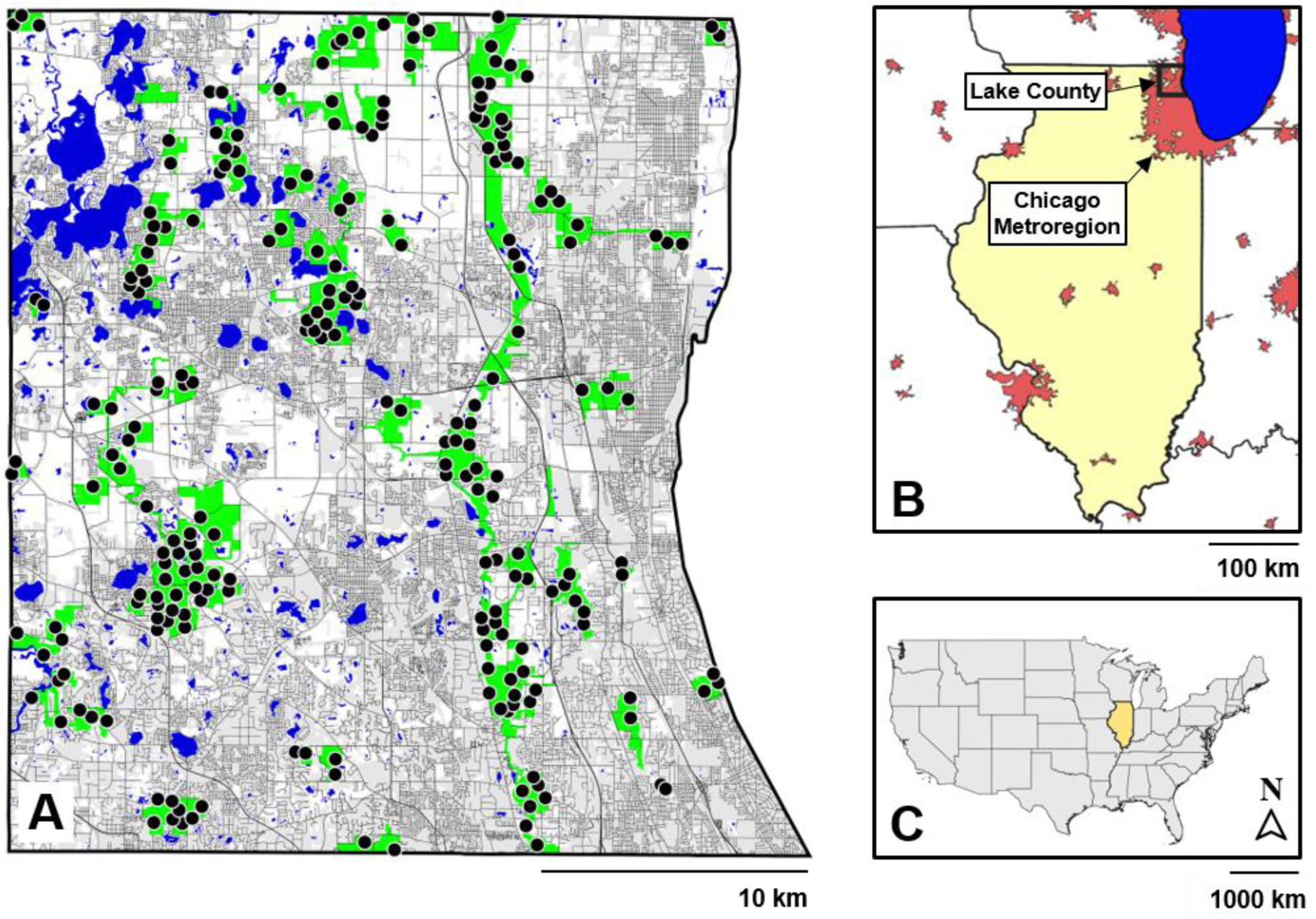
Lake County is a highly urbanized suburb of Chicago, IL, USA. (A) Camera trap locations (black circles, n = 232) and preserves (green polygons, n = 55) monitored from 2009-2018 for mesopredators, along with roads (dark grey lines), open water (blue polygons), and urban land-cover classes (light grey polygons) pooled from the from the 2011 National Landcover Database (Homer et al. 2015). White space indicates other landcover classes which consist mostly of agriculture (center) and emergent wetlands (northwest and northeast corners). (B) Location of Lake County within Illinois (beige) and in relation to the third largest metro region in the USA, Chicago (labeled) and other major US Census designated urban areas (maroon). (C) Location of Illinois (beige) within the United States.

### 2.2. Field Methods

We used remote camera traps (Kelly et al. 2012) to monitor mesopredators, including domestic cats, at 232 monitoring points across 55 Lake County preserves from 2009 to 2018 (Figure 1). As part of the long-term multi-taxa wildlife monitoring program (Cassel 2014, Cassel et al. 2019, 2020, Vanek and Glowacki 2019, Vanek et al. 2019) preserves were initially categorized into two groups (priority and non-priority) based on *a priori* restoration and management goals by LCFPD wildlife ecologists. We used a geographic information system (GIS) to randomly distribute points at an average density of 0.5 points/ha in priority preserves and 0.2 points/ha in non-priority preserves with a minimum of 2 points/preserve and a minimum distance of 400 m between points. Not all preserves could be sampled every year due to the scale of the study, so we used a staggered-entry design starting in 2010 with 18 preserves and 82 points (Table 1). We sampled priority preserves (n = 26) every other year and non-priority preserves (n = 29) every four years. Each year we sampled scheduled preserves (and all monitoring points within) for 4 nights (5 calendar days) over an 8-week period from mid-August through early November. These rapid biodiversity surveys took place in the autumn to maximize the detection probability of native mesopredators as offspring mature and disperse.

**Table 1.** Camera trap effort used to sample mesopredators at 232 monitoring points across 55 preserves from 2010–2018 in Lake County, IL. Preserves were sampled using a staggered-entry design entry and preserves were sampled either every two or four years depending on their priority status. One camera was assigned to each monitoring point and set for 4 trap nights (Monday– Friday; 5 calendar days) during each sampling period. The Trap Days column reflects missing sampling periods due to malfunctioning cameras, theft, etc.

| Year | Preserves<br>Sampled | Points<br>Sampled | Trap<br>Nights | Trap<br>Days |
| --- | --- | --- | --- | --- |
| 2010 | 20 | 105 | 420 | 515 |
| 2011 | 21 | 94 | 376 | 455 |
| 2012 | 21 | 108 | 432 | 540 |
| 2013 | 20 | 87 | 348 | 435 |
| 2014 | 20 | 105 | 420 | 445 |
| 2015 | 20 | 91 | 364 | 455 |
| 2016 | 20 | 106 | 424 | 530 |
| 2017 | 20 | 87 | 348 | 435 |
| 2018 | 20 | 105 | 420 | 525 |
| Total | 55 | 232 | 3552 | 4335 |

During each sampling occasion, we deployed one camera (Leaf River® IR-3BU™, Cuddeback® Ambush™, or Bushnell® Trophy Cam™) within a 100 m buffer of each monitoring point at locations frequented by mesopredators (e.g. game trails or habitat edges). We mounted cameras on trees or metal posts at a height of 0.5 m with a line-of-sight parallel to the ground or slightly downward. We emptied one can of sardines (106 g) on the ground 5 m in front of each camera and cleared any vegetation or debris between the camera and the bait. Cameras were deployed on Mondays and removed on Fridays (thus totaling 4 trap nights over 5 calendar days per sampling occasion). We set cameras to take 1 photo per trigger with a delay of 1 minute. Cameras were checked daily and we replaced bait as needed.

### 2.3. Activity Patterns

We created circular kernel density estimates for free-ranging cats using the *overlap* package in the R Statistical Environment (R Core Team 2018), which uses a *von Mises* kernel density function to accurately represent the circular distribution of time of day (Rowcliffe et al. 2014). We excluded all detections at the same site if they were within 10 minutes of a previous detection to avoid temporal autocorrelation of the same animal triggering a camera repeatedly.

### 2.4. Occupancy Models

We used single-season occupancy models (MacKenzie et al. 2002, Tyre et al. 2003) to investigate the occupancy and detectability of free-ranging cats. This method estimates rates of site occupancy (the probability a site is occupied; *ψ*) and detectability (the probability a species is detected at a site *given a site is occupied*) based on repeated surveys at a site. Estimates of *ψ*explicitly incorporate the uncertainly of detection probabilities < 1. Ignoring the biological reality of imperfect detectability can result in incorrect estimates of wildlife parameter estimates (Gu and Swihart 2004). Occupancy models are useful when surveying large areas because they do not rely on identifying individuals, require only presence-absence data, and allow for both parameters to vary by both site- and survey-specific covariates (MacKenzie et al. 2002, 2017). The alternative for a multi-year study like ours would be to use a dynamic occupancy model (MacKenzie et al. 2003), but we were more interested in site-use and spatial patterns than rates of colonization and extinction. See Bailey & Adams (2005) for an accessible overview of occupancy analysis.

### 2.5. Modeling Procedure

We considered four indices of urbanization we hypothesized would predict occupancy and detectability of free-ranging cats: distance to nearest building, building density, % impervious surface, and area protected by preserves. We developed directional predictions based on how they might influence these parameters (Table 2). The average correlation coefficient between these covariates was |0.48| +- 0.05. We used a GIS to calculate the distance to nearest building, building density, and impervious surface indices using high resolution (1 m) landcover data for Lake County (Chicago Metropolitan Agency for Planning Data Hub 2018). We used a 400 m buffer from the center of each monitoring point for the building density, impervious surface, and preserve area covariates based on the minimum distance between monitoring points, which also corresponds to the recommended buffer distance between houses and areas containing species vulnerable to cat depredation (Hanmer et al. 2017). We used a preserve area covariate instead of a “distance to urban-edge” covariate as used in other studies because it is often arbitrary where the “urban-edge” begins. We included sampling year as a site-specific covariate, along with survey day and temperature as survey-specific covariates to control for any latent heterogeneity in our sampling methodology. We created the temperature covariate using historical data from weather station USC00115961 in Lake County (Midwestern Regional Climate Center 2019).

**Table 2.** Site and survey covariates used to model the occupancy (ψ) and detectability (p) of domestic cats in Lake County, IL preserves from 2010–2018. Hypotheses refer to the predicted relationship between urban covariates and model parameters. Site covariates were used to model occupancy and detectability, while survey covariates can only be used to model detectability. Sampling year was modeled as a categorial covariate.

| Covariate | Parameters and Hypotheses | Median (Range) | Description |
| --- | --- | --- | --- |
| Nearest Building (km) | $\psi$: decrease<br>$p$: decrease | 0.24 (0.08–0.98) | distance from monitoring point to nearest building |
| Building Density (buildings/ha) | $\psi$: increase<br>$p$: increase | 0.22 (0.00–4.66) | number of buildings within 400 m buffer (50.4 ha) around monitoring point |
| Impervious Surface (%) | $\psi$: increase<br>$p$: increase | 3.06 (0.00–24.2) | area impervious surface (roads, pavement, buildings) within 400 m buffer around monitoring point |
| Preserve Area (%) | $\psi$: decrease<br>$p$: decrease | 0.71 (0.15–1.00) | area within 400 m buffer of monitoring point within preserve boundaries |
| Sampling Year | $\psi$: n/a<br>$p$: n/a | N/A (2010–2018) | year monitoring point was sampled |
| Temperature (°C) | $p$: n/a | 14.4 (01.7–28.1) | mean daily temperature of survey period |
| Survey Date | $p$: n/a | 275 (233–309) | ordinal day of year of survey period (275 = 2 Oct) |

We compiled detection histories for each monitoring point-year combination and considered each calendar day a camera was active to be a single survey. Thus, we had a total of 5 survey periods for each point-year combination. We considered each monitoring point-year combination to be a distinct site (i.e. a stacked design) (Fuller et al. 2016, Crum et al. 2017, Goldspiel et al. 2019). We excluded data from the first year of the monitoring program (2009) from our analysis due to low detections for all species and lower effort relative to subsequent years.

We fit models using a maximum likelihood implementation of single-season occupancy analysis within the *unmarked* R package (Fiske and Chandler 2011). This hierarchical model contains two submodels, one for the occupancy component (ecological process; *ψ*), and the other for the detection component (observation process; *p*):

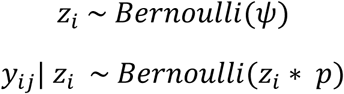

where *z*_*i*_ is a latent variable representing the true occupancy state at site *i*, and *y*_*ij*_ is the observed occupancy status at site *i* during survey *j*, conditional on the true occupancy status *z*_*i*_ (Kéry and Royle 2016). Using a two-stage modeling approach, we first determined the best occupancy sub-model by ranking *a priori* candidate models using a highly parameterized detection sub-model. We then used the most parsimonious occupancy sub-model to determine the best detectability sub-model (MacKenzie et al. 2017). We used the *AICcmodavg* package to rank models with QAIC_c_ (quasi-Akaike’s Information Criterion corrected for small sample sizes) with an overdispersion modifier of 1.1 based on 1000 simulations of the MacKenzie and Bailey Goodness-of-fit Test (MacKenzie and Bailey 2004). We considered models with ΔQAIC_c_ < 2 to have “substantial empirical support” (Burnham and Anderson 2002, Powell and Gale 2015). See the supplemental materials for yearly detection histories, site-specific covariate data, candidate model sets, and R scripts.

### 2.6 Overall Detection Probability

To estimate the number of surveys needed to detect cats during a single 4 trap night, 5 calendar day camera trap survey, we used values from top ranked, detection corrected occupancy model and the formula

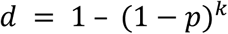

where *p* = the per-survey detection probability and *k* = the number of surveys (Powell and Gale 2015).

### 2.7 Landscape Scale Occupancy

We used the top ranked, detection corrected occupancy model to generate spatially explicit estimates of cat occupancy across all 55 preserves. First, we generated a grid of 25 m x 25 m squares across all preserves using a GIS, then estimated the occupancy value for the centroid of each point using the *predict* function in R. We averaged these predicted occupancy values to estimate mean occupancy for each preserve. In addition, because preserves often consist of distinct geographic units, we also calculated predicted occupancy for each preserve patch, which we defined as each separate polygon in our preserve shapefile layer. For example, a preserve bisected completely by a paved road would consist of two separate patches. In total, we were able to divide the 55 preserves into 159 distinct patches (mean area = 73.1 ha, median = 40.7 ha, SD = 89.5, range = 0.02 – 453.7 ha). We hypothesized that cat occupancy would be higher in smaller preserves and patches (Crooks 2002).

Finally, we compared the estimated occupancy for each preserve against two commonly used landscape/design metrics: preserve size (log ha) and an index of preserve shape (Patton 1975):

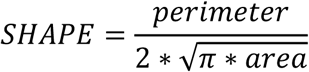

where *perimeter* is the perimeter of a preserve in m and *area* is the area of a preserve in m^2^. A perfectly compact preserve (a circle) would have a SHAPE index of 1, and values larger than 1 indicate an increasingly irregular perimeter to area ratio. We hypothesized that cat occupancy would be lower in more compact preserves and patches (Crooks and Soulé 1999).

## 3. Results

### 3.1 Camera Trapping

We detected cats during all 9 years of the study (Table 3) with the number of detections ranging from four in 2018 to nine in 2012 and 2014. We defined a detection as at least one cat photo at a monitoring point during a survey day. We detected cats 94 times across 45% of preserves (n = 25) and 18% of monitoring points (n = 41). Cats were most often detected only once during a single survey week (n = 37), less frequently twice (n = 10), three times (n = 8), and four times (n = 2). Cats were only detected during all five survey days once. We detected cats at the same monitoring point between years infrequently (n = 11 points) and we tentatively identified at least 16 unique cats at these 11 points (based on a visual assessment). Cats were active during the evening, night, and day with slight peaks of activity before dawn, at noon, and after dusk (Figure 2).

**Table 3.** Domestic cats and species of native terrestrial mesopredators detected in Lake County Forest Preserves via camera traps surveys from 2010–2018. Naïve occupancy is calculated as the number of locations where a species was detected at any point of the nine years divided by the total number of preserves (n = 55) and permanent monitoring points (n = 232).

| Species | Years detected | # photos | Naïve Occupancy |
| --- | --- | --- | --- |

|  |  |  | Preserve | Points |
| --- | --- | --- | --- | --- |
| Raccoon | 9 | 10,197 | 0.98 | 0.92 |
| Opossum | 9 | 13,522 | 0.96 | 0.85 |
| Coyote | 9 | 913 | 0.91 | 0.57 |
| Striped Skunk | 9 | 898 | 0.75 | 0.47 |
| **Domestic Cat** | **9** | **355** | **0.47** | **0.18** |
| Red Fox | 7 | 26 | 0.15 | 0.04 |

**Figure 2.**
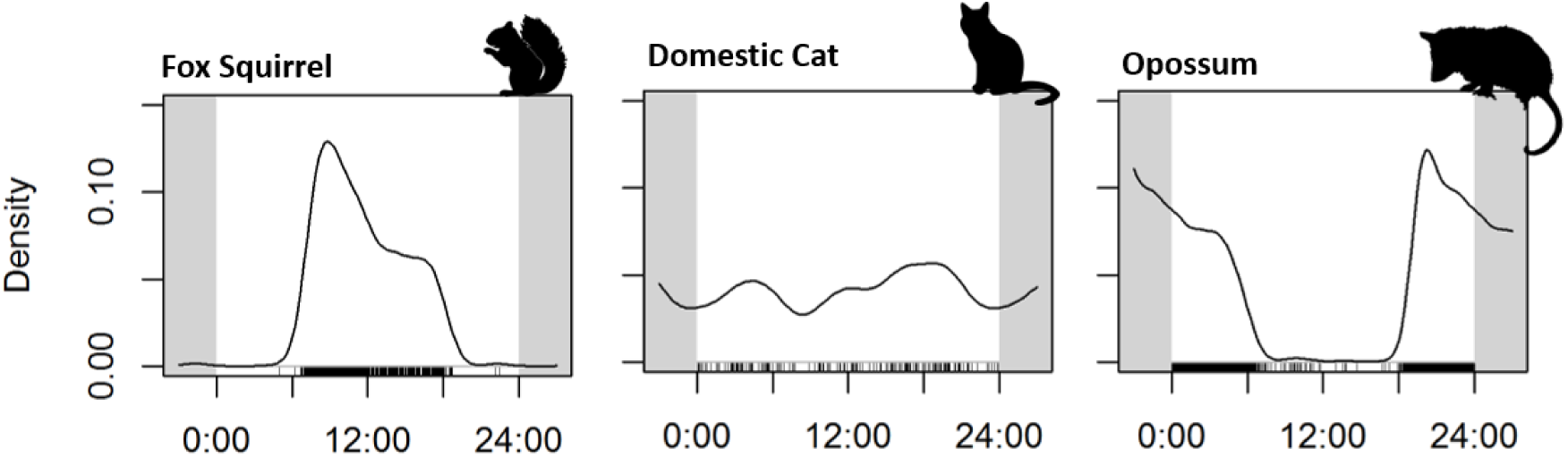
Free-ranging cats (*Felis catus*) in Lake County preserves were active at all times of the day and were not clearly diurnal. Examples of diurnal species, the fox squirrel (*Sciurus niger*), and a nocturnal species, the Virginia opossum (*Didelphis virginiana*), are shown for comparison. Camera trap detections are shown above each x-axis. Activity periods were created using the overlap package in R, which uses a von Mises kernel density function to accurately represent the circular distribution of time of day.

### 3.2. Occupancy and Detection

After removing models (n = 5) with uninformative parameters (Arnold, 2010), we found substantial support (ΔQAIC_c_ < 2) for two occupancy sub-models: *ψ*(IMPERVIOUS SURFACE) and *ψ*(BUILDING DENSITY). These models had very similar levels of support and were essentially equivalent with comparable QAIC_c_ weight and a cumulative model weight of 0.79 (Table 4). These models were 4.1–5.0 times more likely to be the best model than the null model (model likelihood = 0.20), 4.9–6.0 times more likely than the *ψ*(NEAREST BUILDING) model, and 15.1–18.5 times more likely than the *ψ*(PRESERVE AREA) model. There was essentially no support (ΔQAIC_c_ > 12) for models containing the year covariate (Table 4). Full model selection results, including identification of models with uninformative parameters, are presented in the Supplementary Materials.

**Table 4.** Full model set used to evaluate occupancy (ψ) for domestic cats fitted to stacked detection history data from 232 monitoring points across 55 preserves in Lake County, IL from 2010-2018. We modeled occupancy while fixing detection to a sub-global model: p(DOY + TEMP + BDE + DNB + FPA + IMP).

| Model <sup>a</sup> | K <sup>b</sup> | $\Delta\text{QAIC}_c$ <sup>c</sup> | QLL <sup>d</sup> | $w_i$ <sup>e</sup> | Cumulative $w_i$ |
| --- | --- | --- | --- | --- | --- |
| **$p(\text{G}) \psi(\text{IMP})$** | **10** | **0.00** | **-330.97** | **0.21** | **0.21** |
| **$p(\text{G}) \psi(\text{BDE})$** | **10** | **0.40** | **-331.17** | **0.17** | **0.38** |
| $p(\text{G}) \psi(\text{IMP} + \text{FPA})^*$ | 11 | 0.54 | -330.14 | 0.16 | 0.54 |
| $p(\text{G}) \psi(\text{BDE} + \text{FPA})^*$ | 11 | 1.20 | -330.47 | 0.12 | 0.66 |
| $p(\text{G}) \psi(\text{BDE} + \text{IMP})^*$ | 11 | 1.59 | -330.67 | 0.09 | 0.75 |
| $p(\text{G}) \psi(\text{BPR} + \text{IMP})^*$ | 11 | 2.08 | -330.91 | 0.07 | 0.83 |
| $p(\text{G}) \psi(\text{BDE} + \text{BPR})^*$ | 11 | 2.20 | -330.97 | 0.07 | 0.90 |
| $p(\text{G}) \psi(\cdot)$ | 9 | 3.22 | -333.68 | 0.04 | 0.94 |
| $p(\text{G}) \psi(\text{BPR})$ | 10 | 3.59 | -332.77 | 0.04 | 0.98 |
| $p(\text{G}) \psi(\text{BPR} + \text{FPA})$ | 11 | 5.71 | -332.73 | 0.01 | 0.99 |
| $p(G) \psi(FPA)$ | 10 | 5.84 | -333.89 | 0.01 | 1.00 |
| $p(G) \psi(YEAR + IMP)$ | 18 | 12.43 | -328.08 | 0.00 | 1.00 |
| $p(G) \psi(YEAR + BPE)$ | 18 | 13.26 | -328.50 | 0.00 | 1.00 |
| $p(G) \psi(YEAR)$ | 17 | 15.46 | -330.77 | 0.00 | 1.00 |
| $p(G) \psi(YEAR + BPR)$ | 18 | 15.57 | -329.65 | 0.00 | 1.00 |
| $p(G) \psi(YEAR + FPA)$ | 18 | 17.81 | -330.77 | 0.00 | 1.00 |

<sup>a</sup> G = sub-global model; DNB = distance to nearest building; FPA = preserve area; IMP = impervious surface; BDE = building density; (.) = null model (no covariates); \* = model with uninformative parameter(s).
<sup>b</sup> number of model parameters.
<sup>c</sup> difference in quasi-Akaike's Information Criterion corrected for small sample sizes between current model and the top model.
<sup>d</sup> quasi-Log Likelihood
<sup>e</sup> model weight

Because there was a similar level of support for two occupancy covariates, we assessed detection sub-models containing both the building density and impervious surface occupancy covariates. Of these 16 models, only one model was competitive (QAIC_c_ < 2): *p*(PRESERVE AREA) *ψ*(BUILDING DENSITY) (Table 5). This model was 4.4 times more likely to be the best model relative to the second highest ranked model *p*(PRESERVE AREA) *ψ*(IMPERVIOUS SURFACE), and > 600 times more likely than the null detection model. There was essentially no support (ΔQAIC_c_ > 10) for detection models containing other urban covariates, the year covariate, or survey-specific covariates (e.g. temperature, survey day) (Table 5). Full model selection results are presented in the Supplementary Materials.

**Table 5.** Full model set used to evaluate detectability (p) for domestic cats fitted to stacked detection history data from 232 monitoring points across 55 preserves in Lake County, IL from 2010-2018. We modeled detectability using the two top-ranked occupancy submodels, Ψ(BDE) and Ψ(IMP). The top ranked model is bolded.

| Model <sup>a</sup> | K <sup>b</sup> | $\Delta QAIC_c$ <sup>c</sup> | -2QLL <sup>d</sup> | $w_i$ <sup>e</sup> | Cumulative $w_i$ |
| --- | --- | --- | --- | --- | --- |
| **$\psi(BDE) p(FPA)$** | **5.00** | **0.00** | **-334.41** | **0.81** | **0.81** |
| $\psi(IMP) p(FPA)$ | 5.00 | 2.98 | -335.90 | 0.18 | 0.99 |
| $\psi(BDE) p(BPR)$ | 5.00 | 10.91 | -339.87 | 0.00 | 0.99 |
| $\psi(BDE) p(.)$ | 4.00 | 13.03 | -341.97 | 0.00 | 1.00 |
| $\psi(BDE) p(BDE)$ | 5.00 | 13.18 | -341.00 | 0.00 | 1.00 |
| $\psi(BDE) p(IMP)$ | 5.00 | 13.93 | -341.38 | 0.00 | 1.00 |
| $\psi(IMP) p(BPR)$ | 5.00 | 14.18 | -341.50 | 0.00 | 1.00 |
| $\psi(BDE) p(TEMP)$ | 5.00 | 14.67 | -341.75 | 0.00 | 1.00 |
| $\psi(\text{BDE}) p(\text{DOY})$ | 5.00 | 15.11 | -341.97 | 0.00 | 1.00 |
| $\psi(\text{BDE}) p(\text{YEAR})$ | 12.00 | 15.38 | -334.53 | 0.00 | 1.00 |
| $\psi(\text{IMP}) p(\text{BDE})$ | 5.00 | 16.00 | -342.41 | 0.00 | 1.00 |
| $\psi(\text{IMP}) p(\cdot)$ | 4.00 | 17.46 | -344.19 | 0.00 | 1.00 |
| $\psi(\text{IMP}) p(\text{YEAR})$ | 12.00 | 17.56 | -335.62 | 0.00 | 1.00 |
| $\psi(\text{IMP}) p(\text{TEMP})$ | 5.00 | 18.96 | -343.89 | 0.00 | 1.00 |
| $\psi(\text{IMP}) p(\text{IMP})$ | 5.00 | 19.13 | -343.98 | 0.00 | 1.00 |
| $\psi(\text{IMP}) p(\text{DOY})$ | 5.00 | 19.48 | -344.15 | 0.00 | 1.00 |

<sup>a</sup> DNB = distance to nearest building; FPA = preserve area; YEAR = sampling year; IMP = impervious surface; TEMP = mean temperature °C on day of survey; DOY = ordinal day of year of survey; (·) = null model (no covariates).
<sup>b</sup> number of model parameters.
<sup>c</sup> difference in quasi-Akaike's Information Criterion corrected for small sample size between current model and the top model.
<sup>d</sup> quasi-Log Likelihood
<sup>e</sup> model weight

Based on our final detection-corrected model *p*(PRESERVE AREA) *ψ*(BUILDING DENSITY), predicted detection probability decreased with increasing preserve area within the 400 m buffer (*β* = −3.79 ± 0.85 SE) and predicted occupancy increased with the number of buildings within the 400 m buffer (*β* = 0.38 ± 0.15 SE) (Figure 3). Beta values are on the logit scale.

**Figure 3.**
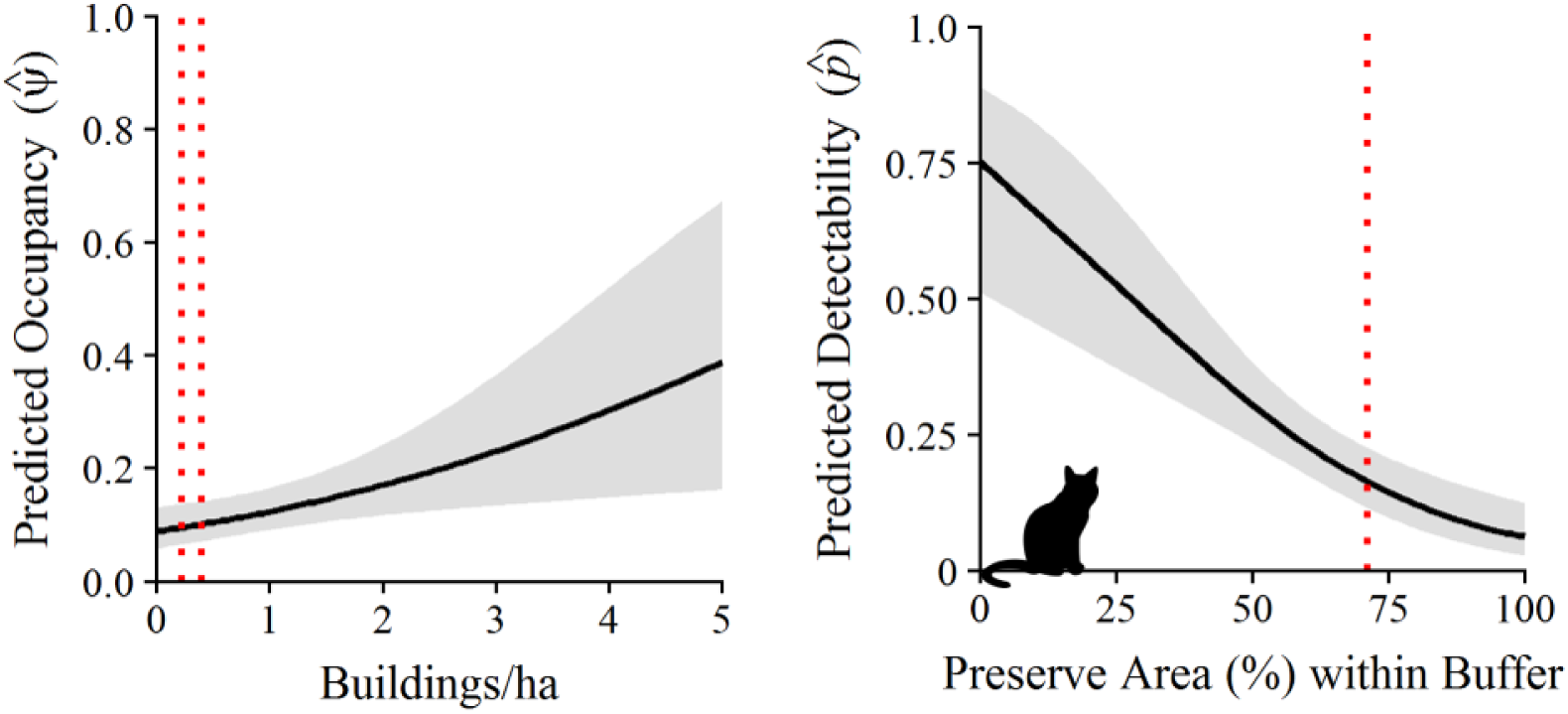
Domestic cat occupancy increased with the number of buildings within a 400 m buffer (50.2 ha) of each monitoring point. The dotted lines represent the median building density (buildings/ha) across all 232 monitoring points (building density _point_ = 0.22) and averaged across all preserves (building density _preserve_ = 0.40). Free-ranging cat detectability decreased with increasing amounts of preserve area within 400 m (50.2 ha) of a monitoring point. The dotted line represents the median preserve area across all 232 monitoring points (preserve area = 71%).

Using values from the final detection-corrected model, we estimated an overall detection probability of 97.4% for a single 4 trap night, 5 calendar day, camera trapping session if the monitoring locations with a 25% preserve area buffer. This overall detection probability drops to 26% at locations that are 100% preserve, and it would take at least four 5-day camera trapping sessions to reach 90% with a single baited camera.

### 3.3. Landscape Scale Patterns

Preserve level building density averaged 0.57 buildings/ha (based on a 400 m buffer around each 25 m x 25 m grid square centroid) and ranged from 0.04–3.68 buildings/ha. Correspondingly, preserve level occupancy was low and averaged 0.11 ± 0.03 SD across all 55 preserves, ranging from 0.09 (95% CI = 0.06 – 0.13) at Mill Creek, Ethel’s Woods, and Gander Mountain to 0.28 (95% CI = 0.15 – 0.48) at Berkeley Prairie (Figure 3). Predicted occupancy across the 159 patches was 27% higher than preserve level occupancy at 0.14 (± 0.07 SD) and ranged from 0.09 (95% CI = 0.06–0.13) to 0.60 (95% CI = 0.20–0.89). As expected, predicated cat occupancy was lower in larger preserves at both the preserve (R^2^ = 0.159, F = 10.02, p = 0.003) and patch level (R^2^ = 0.194, F = 39.2, p < 0.001).

**Figure 3.**
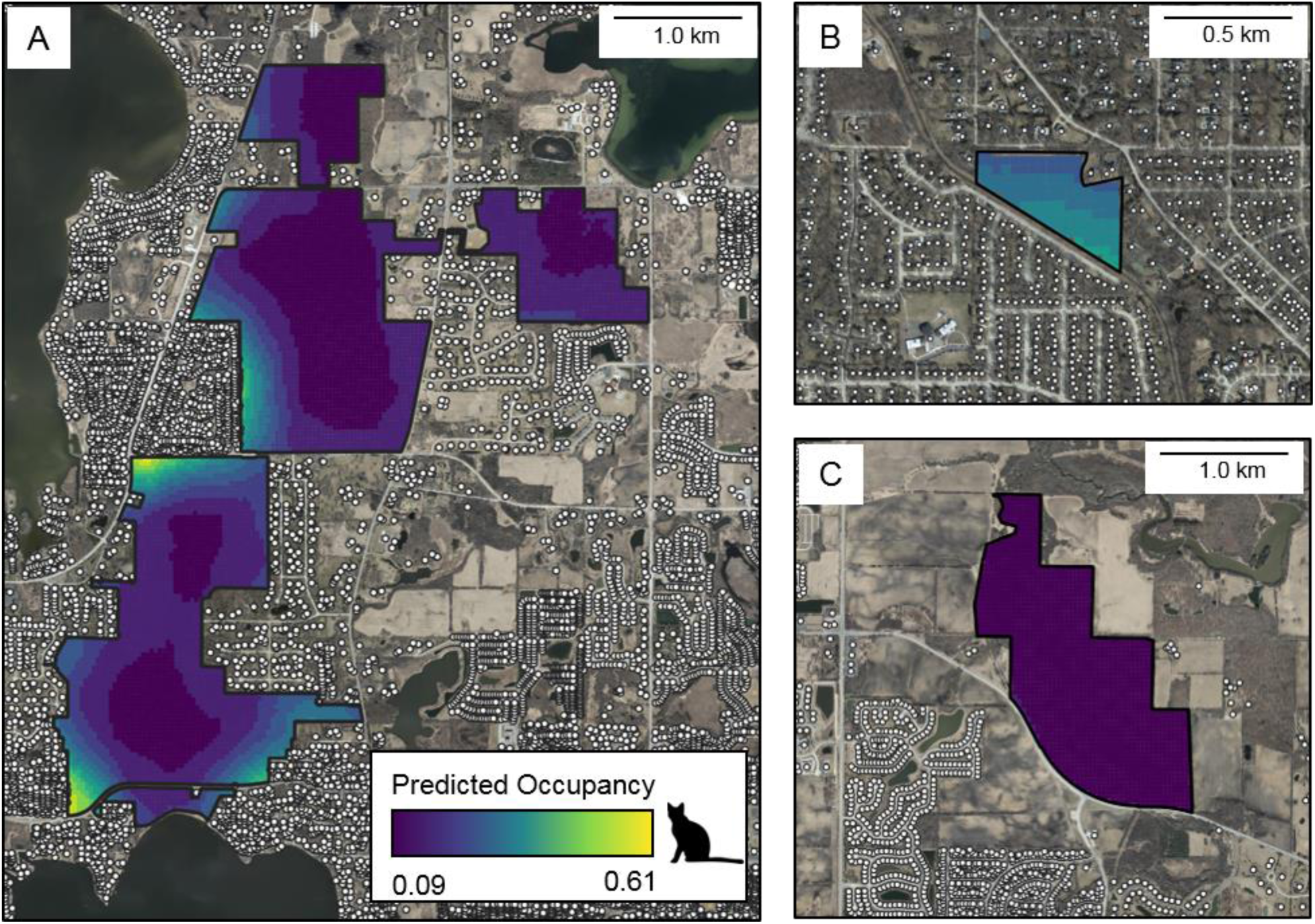
Predicted occupancy for free-ranging cats across three preserves in Lake County, IL. Predicted occupancy at Grant Woods (A) was 0.14 (95% CI 0.91–0.21) and there were pockets of low and high occupancy. Berkeley Prairie (B) was the smallest preserve (7.6 ha) and had the highest predicted occupancy of the 55 preserves at 0.28 (95% CI = 0.15–0.48). C) Mill Creek (112 ha) had the lowest predicted occupancy at 0.09 (95% CI = 0.60–0.13). Building centroids displayed as white circles.

We found no relationship between the SHAPE index and predicted occupancy (Figure 4) at either the preserve (R^2^ = 0.003, F = 0.174, p = 0.673) or patch level (R^2^ = - 0.006, F = 0.004, p = 0.948). Larger preserves tended to be more irregular than smaller preserves (R^2^ = 0.27, F = 19.56, p < 0.0001), but larger patches did not tend to be more irregular than smaller patches (R^2^ = −0.005, F = 0.151, p = 0.698).

**Figure 4.**
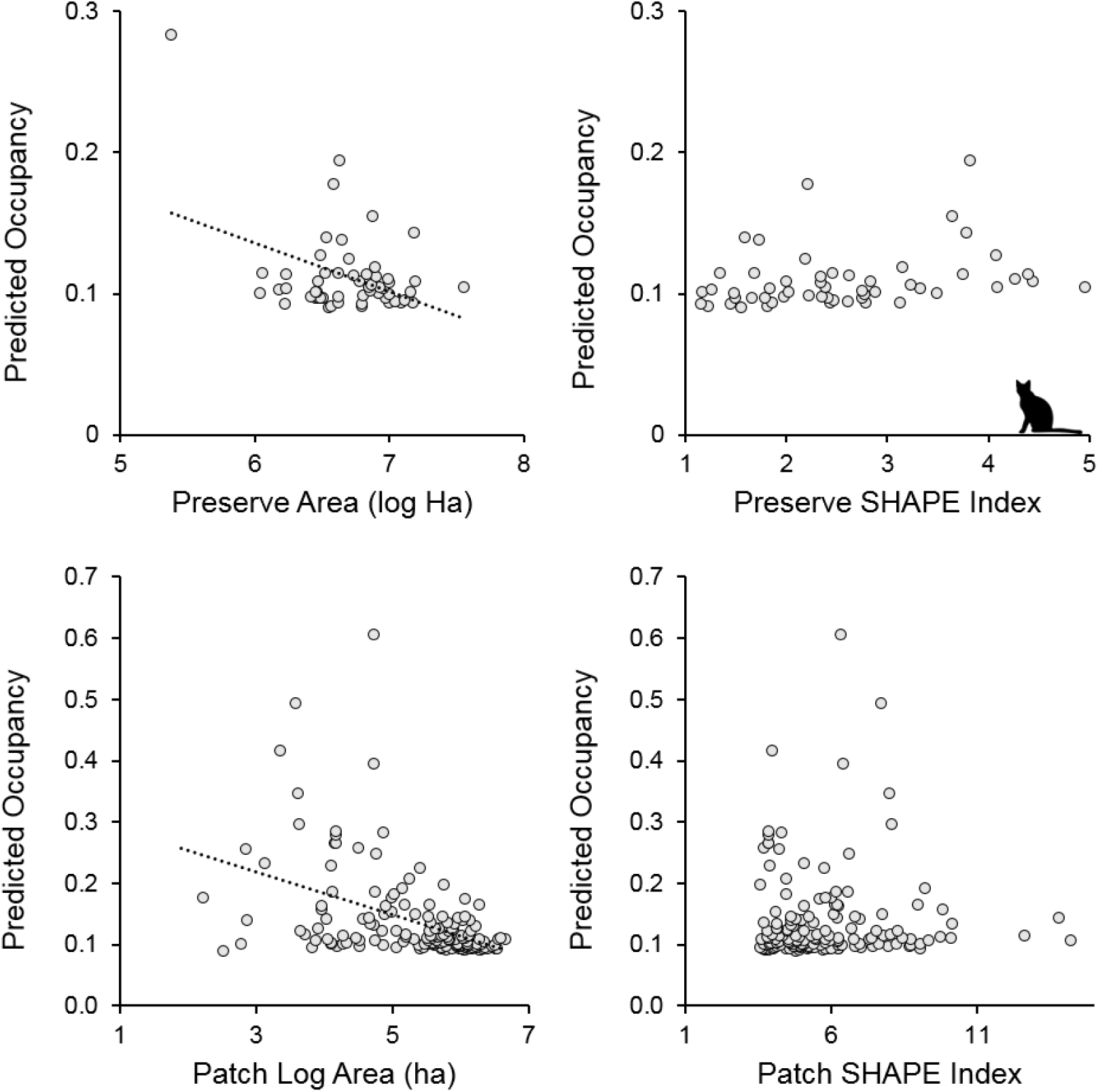
Relationship between predicted occupancy of free-ranging cats, preserve area, and the SHAPE index (a measure of compactness) for 55 sampled preserves in Lake County, IL. A perfectly compact preserve (a circle) would have a SHAPE index of 1, and values larger than 1 indicate an increasingly irregular perimeter to area ratio.

## 4. Discussion

We used occupancy modeling and camera trap data from a long-term wildlife monitoring program to explore the spatial ecology of free-ranging cats within a large network of suburban natural areas. We detected cats in less than half of our preserves and at less than 20% of monitoring points, although cats were likely to be present at more preserves due to the relatively low detection probability across much of the preserve network (see section 4.2 below). In contrast, raccoons, opossums, skunks, and coyotes were widely distributed, and we documented their presence in nearly every preserve, consistent with regional predictions (Gallo et al. 2017, Greenspan et al. 2018). There are few landscape level reports of urban biodiversity (but see Gallo et al. 2017 and Magle et al. 2019), and our work provides important baseline data for land managers and conservation planners.

Cats in our study area were not decidedly nocturnal or crepuscular, but active around the clock (Figure 4). This is consistent with the activity patterns of owned cats (Horn et al. 2011), where feral and unowned cats typically show a nocturnal or strongly crepuscular mesopredators activity pattern (Konecny 1987, Horn et al. 2011, Wang et al. 2015, Cove et al. 2018). Most mesopredators native to the region are crepuscular or nocturnal (Lesmeister et al. 2015). Therefore, if cats are active in preserves during the day, they likely pose an additional risk to diurnal prey species which may not be adapted to diurnal mammalian mesopredators. In addition, free-ranging cats may compete with other diurnal predators, such as raptors (George 1974, Monterroso et al. 2013).

Previous research clearly shows that urban areas can foster large cat populations (Flockhart et al. 2016), and Lake County has a human population density greater than >98% of all counties in the United States. Indeed, with over 240,000 households in Lake County (Planning, Building, and Development Department 2019), there are likely to be over 110,000 individual pet cats, based on the national average of 1.8 cats per household at a cat ownership rate of 25% (American Veterinary Medical Association 2019). We hypothesize that the large population of coyotes in the Chicago Metropolitan Area (Gehrt et al. 2009, 2011) might be limiting cat occupancy of suburban nature preserves in Lake County, as has been proposed in other areas (Crooks and Soulé 1999, Gehrt et al. 2013, Kays et al. 2015). For example, in neighboring Cook County, IL, Gehrt et al. (2013) used GPS collars on sympatric free-ranging cats and coyotes to show habitat partitioning, with cats selecting areas of urban landcover types and avoiding natural areas, which they attribute to predator avoidance. Our results are consistent with this interpretation, with cat occupancy and detection probability lowest in areas further from urban infrastructure.

Free-ranging cat occupancy was influenced by the density of buildings and predicted occupancy more than tripled from points with no nearby buildings to points with 4.5 buildings/ha (Figure 2). This is consistent with the observed patterns of cat activity and suggests that cats are selecting habitat near areas of denser human population, indicative of a population of cats relying on people for subsidy. These results are also consistent with landscape-scale studies linking free-ranging cats to building density. Flockhart et al. (2016) found that cat density increased with building density in Guelph, Ontario, and free-ranging cat occupancy was associated with the density of human-made structures across rural southern Illinois (Morin et al. 2018). Similarly, Krauze-Gryz et al. (2012) linked cat occupancy with distance to nearest building in an agricultural landscape in central Poland. However, our building proximity covariate was poorly ranked relative to building density and even the null model, and the importance of building density was only apparent after modeling detection probability. Moreover, we found relatively little correlation between building density and building proximity (*r* = −0.44). In rural areas, building distance is likely to be important because that building might be the only one near a monitoring location, whereas in urban areas, single buildings are rare, and thus the number of buildings provides more variation for modeling. Care must be taken when selecting indices of urbanization, as indicated by our model selection results and the lack of a strong correlation between our urban covariates.

In contrast to our occupancy results, we found building density to be a poor predictor of detection probability, ranking below the null model. Rather, the amount of preserve within the monitoring area was the top-ranked detection model. That is, detection probability was highest in areas with a greater proportion of non-preserve land (e.g. residential neighborhoods or farm fields) increasing from less than 20% detection probability at interior portions of preserves to more than 75% at the edges of preserves (Figure 3). These results are consistent with our occupancy results, as both building density and the preserve area around a monitoring point are associated with preserve boundaries. In an urban reserve in New Zealand, Woolley and Hartley (2019) found that detection rates at cameras near the preserve edge were 6 times greater than at cameras just 100 m into the preserve. Similarly, in a suburban preserve in New York, Kays and DeWan (2004) found that free-ranging cats were rarely detected at scent stations > 50 m from the neighborhood/preserve edge. As detection probability can be influenced by abundance (Royle and Nichols 2003), our occupancy and detection results strongly suggest that cat populations are highest near the edges of suburban natural areas. Therefore, species vulnerable to cat predation or competition along urban-natural edges are likely to be at higher risk in these areas.

Our use of rapid biodiversity surveys was effective at detecting cats at areas near preserve edges. In contrast, much greater survey effort was needed to detect cats at the interior portions of most preserves (i.e. where occupancy is low). While a single week (5 calendar days) of camera trapping was effective for detecting cats near preserve edges, it would take more than a month to reach a 90% detection probability with a single baited camera. This echoes recent recommendations that up to 4 weeks of camera trapping may be needed to obtain precise estimates of local detection probabilities (Kays et al. 2020). Thus, free-ranging cats might go un-noticed in rapid biodiversity surveys in larger urban natural areas. We suggest more than one camera be used in rapid biodiversity surveys of large urban preserves.

Predicted occupancy across the preserve network was low and averaged less than 12%. Most areas of higher occupancy were located near the boundaries of preserves. For example, one of our largest sampled preserves, Grant Woods, had areas of high predicted occupancy where preserve boundaries bordered densely populated neighborhoods. At the same time, large portions of the preserve had low levels of occupancy, as building density in these areas was near or at zero (Figure 4). Our predicted occupancy value is remarkably similar to the predicted occupancy of free-ranging cats from a mosaic of public and private land across 16 counties in rural southern Illinois, where cat occupancy was higher on private land and also linked to anthropogenic features (structures/ha) (Morin et al. 2018). Thus, cats are likely to be present across a small but consistent proportion of natural areas in the Midwestern United States, with pockets of higher occupancy associated with built-up areas.

We found a negative relationship between preserve/patch size and predicted occupancy. This is consistent with our hypothesis and similar to the findings of Crooks (2002). However, the relationship was weak, explaining no more than 20% of the variance at either scale. For example, while the smallest preserve, Berkeley Prairie, had the highest levels of predicted occupancy, it was an outlier in terms of size, at a fourth the size of the next smallest preserve and 20 times smaller than the median preserve. In addition, the predicted occupancy was 1.5 times higher than the preserve with the second highest predicted occupancy. Further, removing Berkeley Prairie from the analysis renders the preserve-scale regression non-significant. There were no obvious outliers at the patch scale. Except for extremely small patches and preserves, we suggest that size alone is a poor predictor of free-ranging cat occupancy in suburban nature areas, as cat occupancy can still be high in portions of larger preserves adjacent to dense residential areas (Figure 4).

Contrary to our predictions, we found no relationship between the SHAPE index and predicted cat occupancy at either the preserve or patch scale. That is, compact preserves had similar levels of occupancy as to more irregularly shaped preserves. Crooks and Soulé (1999) found that smaller habitat fragments had higher cat abundance, which they attributed to smaller patches having “proportionately more urban edge and therefore greater access by housecats bordering the fragment.” However, that study examined mostly small linear fragments (mean area = 13.8 ha), and only one of the 37 fragments was larger than 100 ha (Soule et al. 1988). In contrast, 73% of the preserves in our study were greater than 100 ha, and our largest preserve, Lakewood had the highest SHAPE index, equivalent to 400% more edge than a circular preserve of the same area. This exemplifies the reality of urban preserves, which are rarely designed, but rather are often obtained and expanded opportunistically. For example, since its inception in 1968, Lakewood has more than doubled in size through 39 individual acquisitions (XXX, unpublished data, masked for double-blind peer review). As with preserve size, highly irregular preserves and patches can occur in both areas of high and low building density, even in urban areas. Thus, nearby building density should be the primary concern for suburban land managers, not proxies like preserve size or irregularity.

## 5. Conclusions

Free-ranging cats are a threat to biodiversity, but we have a limited understanding of their ecology and distribution in urban preserves. Our results show that free-ranging cats occur sporadically throughout nature preserves in Lake County. However, cat occupancy was low relative to native mesopredators, possibly due to high coyote occupancy. Cats were active during all times of the day and night, whereas native mesopredators were mostly nocturnal. Overall, these results suggest that most free-ranging cats within the preserves were not feral (e.g. living independent of humans) but were more likely pet cats with access to the outdoors. This has important implications for the management of free-ranging cats in Lake County, as the control of free-ranging cats is a contentious issue (Ash and Adams 2003, Longcore et al. 2009, Loyd and Miller 2010*a, b*, McDonald et al. 2015, Loss and Marra 2018, Loss et al. 2018, Woolley and Hartley 2019).

Traditional measures of preserve design (i.e. shape and size) may not accurately predict the risk of free-ranging cats. We suggest that urban land managers interested in the conservation and reintroduction of cat-sensitive species to urban natural areas consider the surrounding urban matrix in their decision-making process. In addition, urban ecologists should consider multiple indices of urbanization in their analyses instead of assuming all urban metrics are all equivalent. Finally, while the cat occupancy may be low in urban nature preserves, we caution against complacency as even low numbers of cats can cause substantial harm to biodiversity and human health.

